# Long-lasting effects of proanthocyanidins in rat prevent cafeteria diet effects by limiting adipose tissue accrual: Long-lasting effects of GSPE limiting WAT accrual

**DOI:** 10.1101/570994

**Authors:** Iris Gines, Katherine Gil-Cardoso, Joan Serrano, Àngela Casanova-Marti, Maria Lobato, Ximena Terra, M Teresa Blay, Anna Ardevol, Montserrat Pinent

**Author notes:** Corresponding author: Anna Ardévol, Universitat Rovira i Virgili, Departament de Bioquímica i Biotecnologia, c/ Marcel·lí Domingo nº1, 43007 Tarragona, Spain.

## Abstract

A dose of proanthocyanidins with satiating properties proved to be able to limit body weight increase several weeks after administration under exposure to a cafeteria diet. Here we describe the molecular targets and the duration of the effects. We treated rats with 500 mg GSPE/kg BW for ten days. Seven or seventeen weeks after the last GSPE dose, while animals were on cafeteria diet, we used RT-PCR to measure the mRNA of the key energy metabolism enzymes from liver, adipose depots and muscle. We found that a reduction in the expression of adipose Lpl might explain the lower amount of adipose tissue in rats seven weeks after the last GSPE dose. Liver showed increased expression of Cpt1a and Hmgs2 together with a reduction in Fasn and Dgat2. In addition, fatty oxidation (Oxct1 and Cpt1b mRNA) was increased in muscle. However, after seventeen weeks, there was a completely different gene expression pattern. As conclusion, seven weeks after the last GSPE administration there was a limitation in adipose accrual that might be mediated by an inhibition of the gene expression of the adipose tissue Lpl. Concomitantly there was an increase in fatty acid oxidation in liver and muscle.

## Introduction

Excessive adipose tissue significantly increases the risk and prognosis of metabolic syndrome (diabetes mellitus type 2, cardiovascular disease, hyperlipidemia, nonalcoholic fatty liver disease) and several types of cancer.^1^ The causes of excessive body weight are diverse, one of the most prevalent in developing and developed countries being excessive or bad quality food intake^2^.

Proanthocyanidins (PACs) are a group of polyphenols that are widespread in nature (in fruits, vegetables and their beverage products). They have been described as bioactive agents against several unhealthy situations. More specifically, they have the well-documented effect of limiting lipid accumulation and favouring lipid oxidation in the organism^3^ Their effect as specific inhibitors of fat digestion^4^ and absorption.^5^ Furthermore, PACs favour lower RQ(Respiratory Quotient) ^6,7^, due to a higher fat oxidation in liver and muscle^6^. As they are a group of different compounds, some of the effects could be explained by their interaction with molecules or structures located in the gastrointestinal lumen^8,6^. They protect against cafeteria diet-induced damage to the intestinal barrier and are anti-inflammatory agents.^9^ However, the absorbable low-molecular weight flavanols reach intracellular targets inside the body, where they act on different molecular targets to induce increased energy expenditure,^3^ and prevent cholesterol increase in the organism^10^, acting as antihipertensives^10^, antioxidants^11^) and maintaining glucose homeostasis^12^.

The diversity of structures in proanthocyanidin-rich extracts and their interactions are the reason why some of these effects are highly dependent on the dose used for the studies and the physiological state of the animal ^7^. Most studies prove that PACs correct cafeteria diet-induced damage^13,14^. Some studies focus on their possible preventive role in obesity-related pathologies (that is, when they are administered from the beginning of obesogenic diets) ^15^. However, only very few studies have analysed their long-term effects after sub-chronic treatment^16^, ^17^. We have recently attempted to determine the best moment to administer GSPE (Grape seed proanthocyanidin extract) (500 mg/kg) so that it acts most effectively against the damaging effects of an obesogenic diet such as the cafeteria diet^17^. The results showed that PACs had a surprisingly long-lasting effect on body weight that needed a more in-depth analysis. In the present study, we have further analysed the long-lasting effects of sub-chronic GSPE treatment. We compare the duration of their effects, mainly on the energy metabolism and adipose depots, 7 weeks or 17 weeks after the last GSPE dose.

## Materials and methods

### Proanthocyanidin extract

The grape seed extract enriched in proanthocyanidins (GSPE) was kindly provided by *Les Dérivés Résiniques et Terpéniques* (Dax, France). According to the manufacturer, the GSPE used in this study (Batch number: 124029) contains monomers of flavan-3-ols (21.3%), and dimers (17.4%), trimers (16.3%), tetramers (13.3%) and oligomers (5-13 units; 31.7%) of proanthocyanidins. A detailed analysis of the monomeric to trimeric structures can be found in the work by Margalef and col.^18^

### Animal treatments

Female rats (Harlan Rcc:Han), each weighing 240-270 g, were purchased from Charles River Laboratories (Barcelona, Spain). After one week of adaptation, they were individually caged in the animal quarters at 22°C with a 12-hour light/12-hour dark cycle and fed *ad libitum* with a standard chow diet (Panlab 04, Barcelona, Spain) and tap water. Experiments were performed after a period of acclimation.

#### Short cafeteria (SC) experiment

The animals were randomly distributed into two experimental groups (n=7) and fed a standard chow diet *ad libitum* (figure 1, supplementary materials). One group of animals received 500 mg GSPE/Kg bw together with a simplified high-fat-high-sucrose diet for 10 days. The diet consisted of a palatable hypercaloric emulsion presented in an independent bottle, containing (by weight) 10% powdered skimmed milk, 40% sucrose, 4% lard and 0.35% xanthan gum as a stabilizer.^19^ The GSPE dissolved in tap water was orally gavaged to the animals at 18:00 in a volume of 500 µL, one hour after all the available food had been removed. The animals not supplemented with GSPE received water as a vehicle. After 10 days of treatment, all the animals were kept for 18 days on a standard chow diet. Afterwards, the rats started with **the cafeteria diet challenge for 35 days (SC)**. The cafeteria diet consisted of standard chow, bacon, carrots, and sugared milk, which induces voluntary hyperphagia.^20^ This diet was offered *ad libitum* every day.

**Figure 1.**
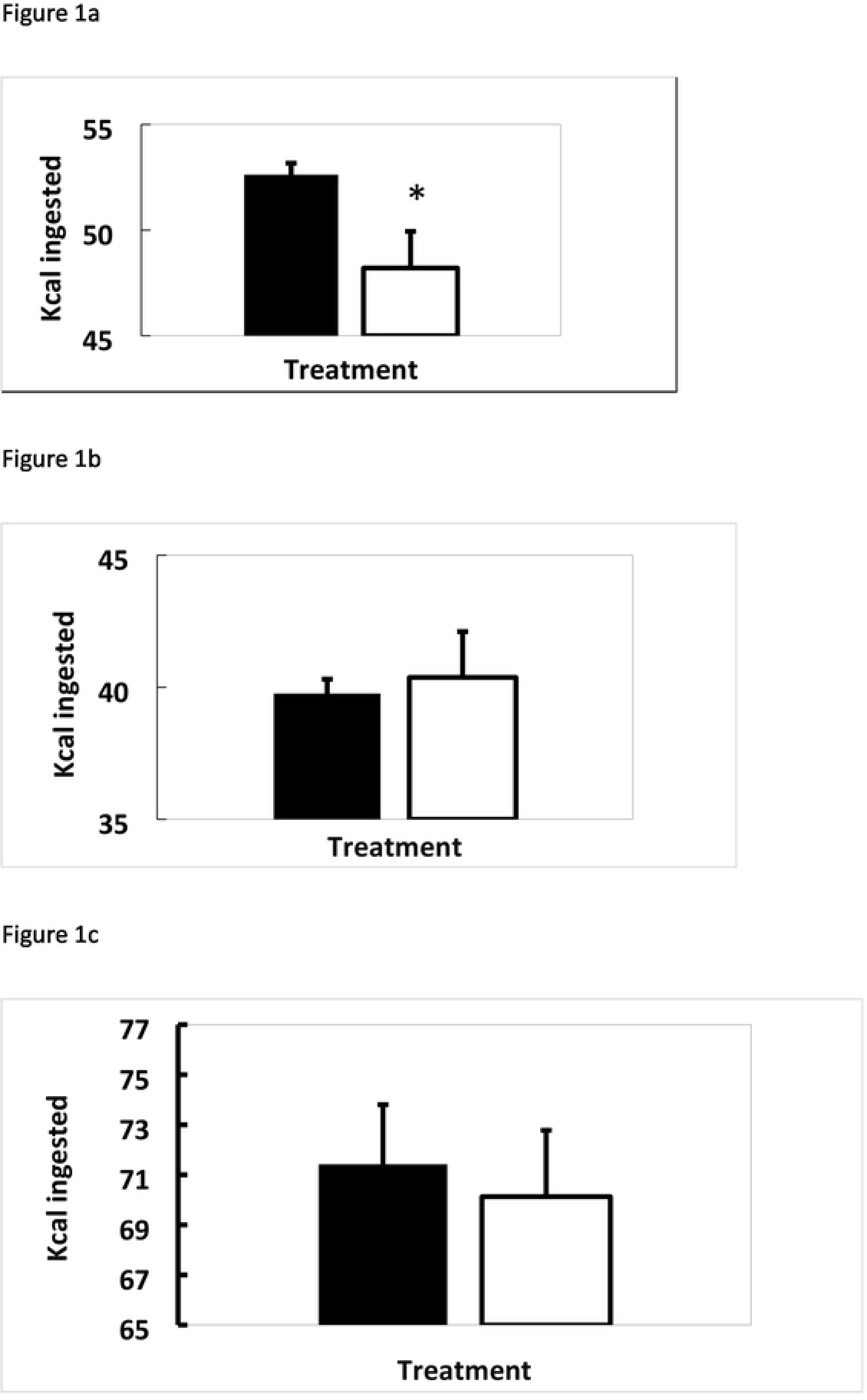
Food intake in the short-challenge group in different diet periods. Food intake was measured 20 hours after the daily food had been replaced for each diet administered. The black column indicates animals not treated with GSPE. The white column indicates animals treated with 0.5 g/Kg BW of GSPE for the first 10 days of treatment. The results showed the mean data obtained from each measurement throughout the period. a) Mean food intake from the daily measurement for the first ten days of treatment with a tasty diet. b) Mean food intake from measurements taken during the eighteen days of treatment with a chow diet. c) Mean food intake from measurements during the thirty-five days with a cafeteria diet. Statistical differences identified by T-test are defined by * when p<0.05 between treatments

#### Long cafeteria (LC) experiment

A second group of thirty female Wistar rats was challenged with a long-term cafeteria treatment (LC), which was initially similar to the treatment described above. They were organized in three experimental groups (n=10), (figure 1, supplementary materials). One group was given the same amount of GSPE every day for 10 days at the same time, and the control group received the same amount of tap water. During the GSPE treatment period, all the rats were fed standard chow diet. On day eleven, a standard group (STD) stayed on the chow diet, and the two remaining groups started a cafeteria challenge, which in this case was maintained for 17 weeks.

In both experiments, the GSPE treatments were intragastrically (i.g.) administered 1 h before the onset of the dark cycle. Food intake was measured 20 hours after the daily chow replacement with an accuracy of 0.01 g.

### Blood and Tissue Collection

At the end of the study, the rats were fasted for 1-4 hours, anesthetized with sodium pentobarbital (70 mg/kg body weight; Fagron Iberica, Barcelona, Spain) and exsanguinated from the abdominal aorta. The blood was collected using heparin (Deltalab, Barcelona, Spain) as an anticoagulant. Plasma was obtained by centrifugation (1500g, 15 minutes, 4°C) and stored at −80°C until analysis. The different white adipose tissue depots (retroperitoneal (rWAT), mesenteric (mWAT) and periovaric (oWAT)), brown adipose tissue (BAT), liver and pancreas were rapidly removed and weighed.

All the procedures were approved by the Experimental Animal Ethics Committee of the Universitat Rovira i Virgili (code: 0152S/4655/2015).

### Plasma metabolites and hormones

Plasma β-hydroxybutyrate was analysed by colorimetry (BEN, Mainz, Alemania). Total ghrelin from plasma samples was analysed with an extraction-free total ghrelin enzyme immunoassay (Phoenix Pharmaceuticals, Burlingame, USA). ^21^. Plasma insulin and glucagon levels were analysed with rat ELISA kits (Mercodia, Sweeden). Plasma leptin levels were analysed with an ELISA kit (Millipore, Billerica, MA, USA).

### Tissue triglycerides and mRNA quantification

Pancreatic triglycerides were assayed according to Castell et al.^22^ Total RNA was extracted using Trizol (Ambion, USA) and trichloromethane-ethanol (Panreac, Barcelona, Spain) and purified using an RNA extraction kit (Qiagen, Hilden, Germany). Complementary DNA was obtained using the High Capacity cDNA Reverse Transcription Kit (Applied Biosystems, Madrid, Spain), and the quantitative reverse transcriptase-polymerase chain reaction (qRT-PCR) amplification was performed using TaqMan Universal PCR Master Mix and the respective specific TaqMan probes (Applied Biosystems, Madrid, Spain). The relative expression of each mRNA was calculated against the control group using the 2^−ΔΔCt^ method, with cyclophilin A as reference.

### Statistical analysis

The data are represented as the mean ± standard error of the mean (SEM). Statistical comparisons between groups were assessed by the T test. Analyses were performed with XLStat 2017.01 (Addinsoft, Spain). P-values <0.05 were considered statistically significant.

## Results

### Sub-chronic treatment with GSPE reduces food intake in rats on a palatable diet

In previous studies we used a GSPE dose that has satiating properties in animals on a chow diet.^7,23^ Here, we reproduce this effect in animals with an enhanced appetite because they were offered a tasty diet. **Figure 1a** shows a 10% reduction in the total food intake of the healthy females while they were treated with GSPE. **Table 1** shows that this reduction was due to the amount of hypercaloric emulsion ingested.

**Table 1.** Characteristics of food intake during the short cafeteria study. GSPE was administered for 10 days together with a tasty diet. After the GSPE treatment stopped, the rats were put on an 18-day chow diet and then a 35-day cafeteria diet. *: (P<0.05 vs C; T test)

|  | Cafeteria | GSPE pre-treated rats |
| --- | --- | --- |
| Initial treatment: 10 days, tasty diet |  |  |
| Chow ingested (g) | 8.34 ± 0.38 | 9.10 ± 0.53 |
| Hypercaloric emulsion ingested (g) | 14.01 ± 0.82 | 10.54 ± 1.07 * |
| Carbohydrates (Kcal) | 43.04 ± 0.78 | 39.15 ± 0.60* |
| Lipids (Kcal) | 6.84 ± 0.19 | 5.90 ± 0.21* |
| Protein (Kcal) | 7.07 ± 0.17 | 7.12 ± 0.22 |
| Final treatment: 35-day cafeteria diet |  |  |
| Carbohydrates (Kcal) | 43 ± 4 | 38 ± 2.0 |
| Lipids (Kcal) | 28 ± 1 | 28 ± 0.6 |
| Protein (Kcal) | 8.4 ± 0.3 | 8.0 ± 0.2 |

The effects on food intake disappeared when GSPE administration finished, as previously shown.^7^ **Figure 1b** shows that there was no difference between groups in the kilocalories (Kcal) ingested over the eighteen days after GSPE treatment, when animals received a standard chow diet. During this period, the rats obtained 68% of energy from CH, 12% from lipids and 20% from protein. When the animals were subsequently subjected to a short (35 days) cafeteria diet, the amount of Kcal ingested was not different between the groups either (f**igure 1c)**. During this last period of the study, animals obtained **5**4% ± 0.020 of energy from carbohydrates (CH) (mainly from the sucrose included in the milk), **36**% ± 0.02 from lipids and **11** % ± 0.003 from protein. As mentioned, these percentages were not statistically different for GSPE treated animals (51% ± 0.01; **38**% ± 0.009; **11**% ± 0.001, from CH, lipids and protein, respectively).

### A reduction in the expression of adipose LPL might explain the lower amount of adipose tissue in GSPE pre-treated rats

We have shown that a 10-day pre-treatment with GSPE followed by a cafeteria diet led to a reduction in adiposity and RQ^17^ after 53 days. **Table 2** shows that in the GSPE pre-treated group there is a statistically significant reduction of around 35% in the size of subcutaneous depots (estimated by the difference between total adipose contents measured by RMN and the weighed intraabdominal depots). Although the mRNA levels of lipid metabolism genes did not help to explain it, there was a statistically significant effect on the ratio between *Cpt1b(*carnitine palmitoyltransferase 1b) vs *Fasn(*fatty acid synthase), suggesting a trend towards a higher oxidative profile in the subcutaneous depot. The next most abundant WAT depot is the periovaric WAT, which did not show any differences between the control and GSPE groups in either weight or gene expression. Instead, the retroperitoneal and mesenteric depots were of statistically different sizes due to the GSPE treatment (reductions of 23 and 35%, respectively). However, there were no significant changes in the gene expression profile, only a tendency to present lower *Fasn* mRNA levels in mesenteric WAT (**table 2**).

**Table 2.**
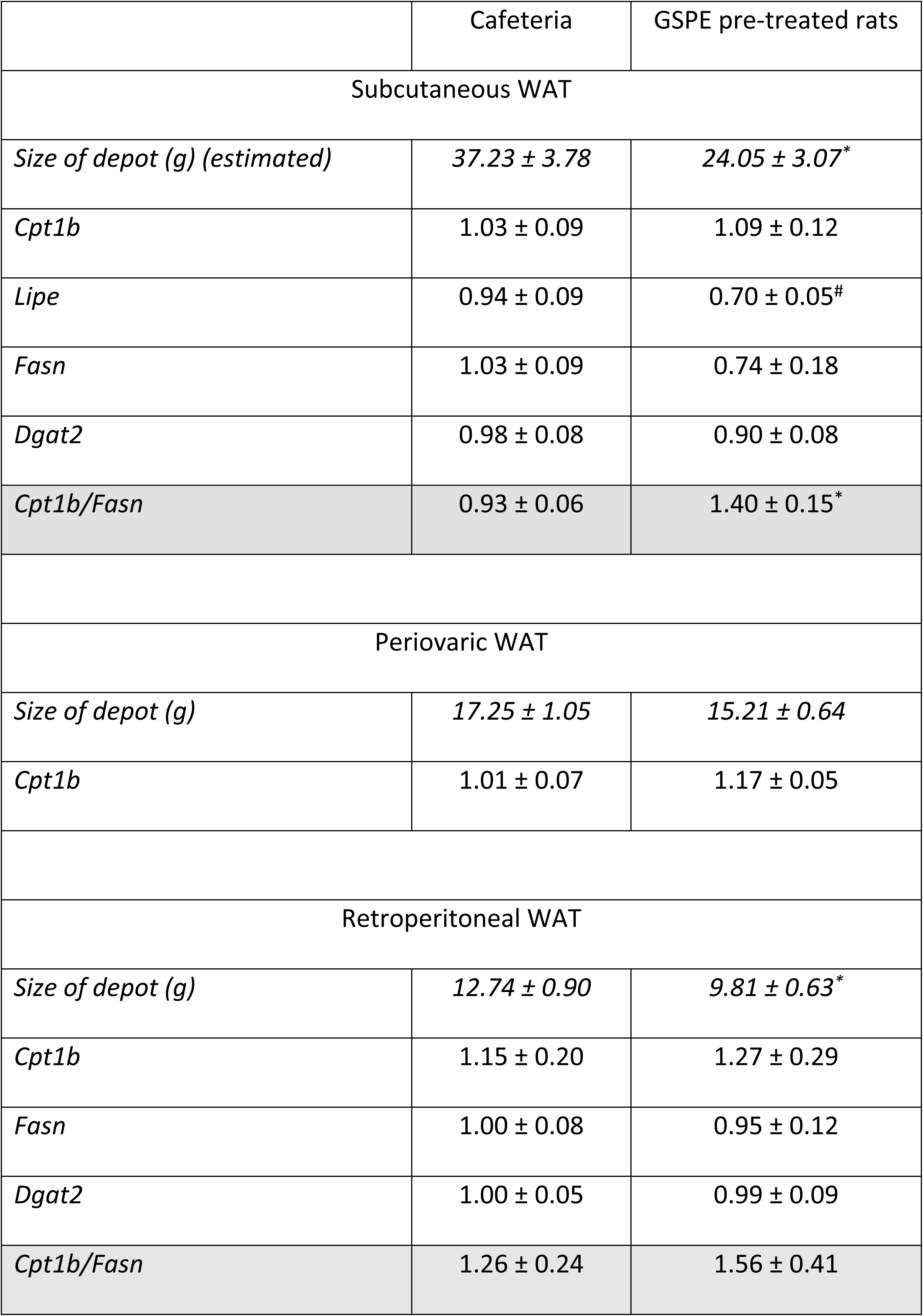
GSPE effects on white adipose depots in the short cafeteria treatment. Adipose depots were obtained at the end of the treatment and each depot was weighed. RT-PCR was used for each gene (results are presented as arbitrary units versus cafeteria group). The T test was used to determine statistical differences highlighted as *p<0.05 vs cafeteria treatment; #p<0.1 vs cafeteria group. Lipe (lipase E, hormone sensitive type)

|  | Cafeteria | GSPE pre-treated rats |
| --- | --- | --- |
| Subcutaneous WAT |  |  |
| *Size of depot (g) (estimated)* | *37.23 ± 3.78* | *24.05 ± 3.07\** |
| *Cpt1b* | *1.03 ± 0.09* | *1.09 ± 0.12* |
| *Lipe* | *0.94 ± 0.09* | *0.70 ± 0.05<sup>#</sup>* |
| *Fasn* | *1.03 ± 0.09* | *0.74 ± 0.18* |
| *Dgat2* | *0.98 ± 0.08* | *0.90 ± 0.08* |
| *Cpt1b/Fasn* | *0.93 ± 0.06* | *1.40 ± 0.15\** |
| Periovaric WAT |  |  |
| *Size of depot (g)* | *17.25 ± 1.05* | *15.21 ± 0.64* |
| *Cpt1b* | *1.01 ± 0.07* | *1.17 ± 0.05* |
| Retroperitoneal WAT |  |  |
| *Size of depot (g)* | *12.74 ± 0.90* | *9.81 ± 0.63\** |
| *Cpt1b* | *1.15 ± 0.20* | *1.27 ± 0.29* |
| *Fasn* | *1.00 ± 0.08* | *0.95 ± 0.12* |
| *Dgat2* | *1.00 ± 0.05* | *0.99 ± 0.09* |
| *Cpt1b/Fasn* | *1.26 ± 0.24* | *1.56 ± 0.41* |

Brown adipose tissue was also analysed but there were no changes due to GSPE either in weight (0.91 ± 0.07 for the cafeteria group; 0.77 ± 0.04 for the GSPE-pre-treated group) or in *Cpt1b* gene expression (1.02 ± 0.0 for the cafeteria group; 1.15 ± 0.2 for the GSPE-pre-treated group) suggesting a lack of effect on oxidative activity in this tissue.

Next, to identify the effects of GSPE on triglyceride entry into adipose tissue, we measured the gene expression of the genes related to this process: *Lpl*(lipoprotein lipase), the enzyme that hydrolyses triglycerides into fatty acids and glycerol, before their uptake into the cell; *Cd36*(CD36 molecule), involved in free fatty acid uptake into the cell; and Aqp7(aquaporin 7), which facilitates the efflux of glycerol from the cell. **Figure 2** shows that the amount of Lpl in the mesenteric depot was highly reduced, as was the amount of Aq7. There were no statistically significant differences for the fatty acid transporter (Cd36).

**Figure 2.**
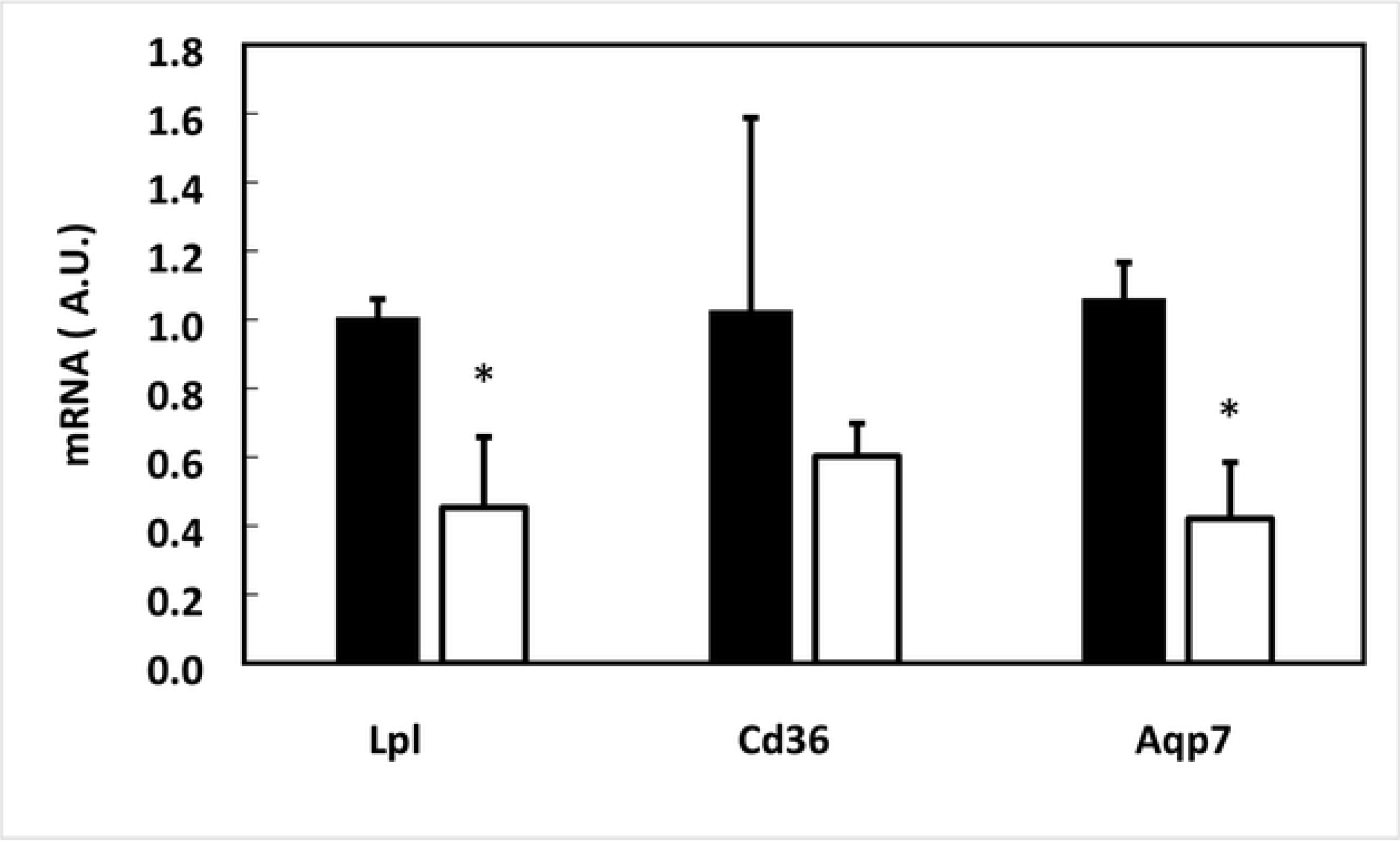
Effect of GSPE pre-treatment on mesenteric adipose gene expression in the short-challenge group at the end of the study. Rats were treated with 0.5 g/Kg BW for the first 10 days, and then they were put on a chow diet for 18 days and a cafeteria diet for 35 days. The black column indicates animals not treated with GSPE. The white column indicates animals treated with 0.5 g/Kg BW of GSPE for the first 10 days of treatment. The mRNA extracted from mesenteric adipose was quantified by RT-PCR and the relative gene expression of Lpl, Cd36 and AqpP7 was obtained by the DDCt method in each gene. The data are the mean ± SEM (n=7). Statistical differences identified by T-test are defined by * when p<0.05 between treatments.

### Lipid oxidation in liver and muscle is higher

In our search for an explanation for the lower RQ observed in GSPE pre-treated rats,^17^ we analysed liver and muscle gene expressions. **Figure 3a** shows a significant increase in *Cpt1a* (carnitine palmitoyltransferase 1a) and *Hmgs2*(3-hydroxy-3-methylglutaryl-CoA synthase 2), suggesting the increased oxidation of fatty acids and the active synthesis of ketone bodies in the liver of GSPE-pre-treated animals. In addition, decreased *Fasn* and *Dgat 2*(diacylglycerol O-acyltransferase 2) expression suggested decreased synthesis and esterification of fatty acids.

**Figure 3.**
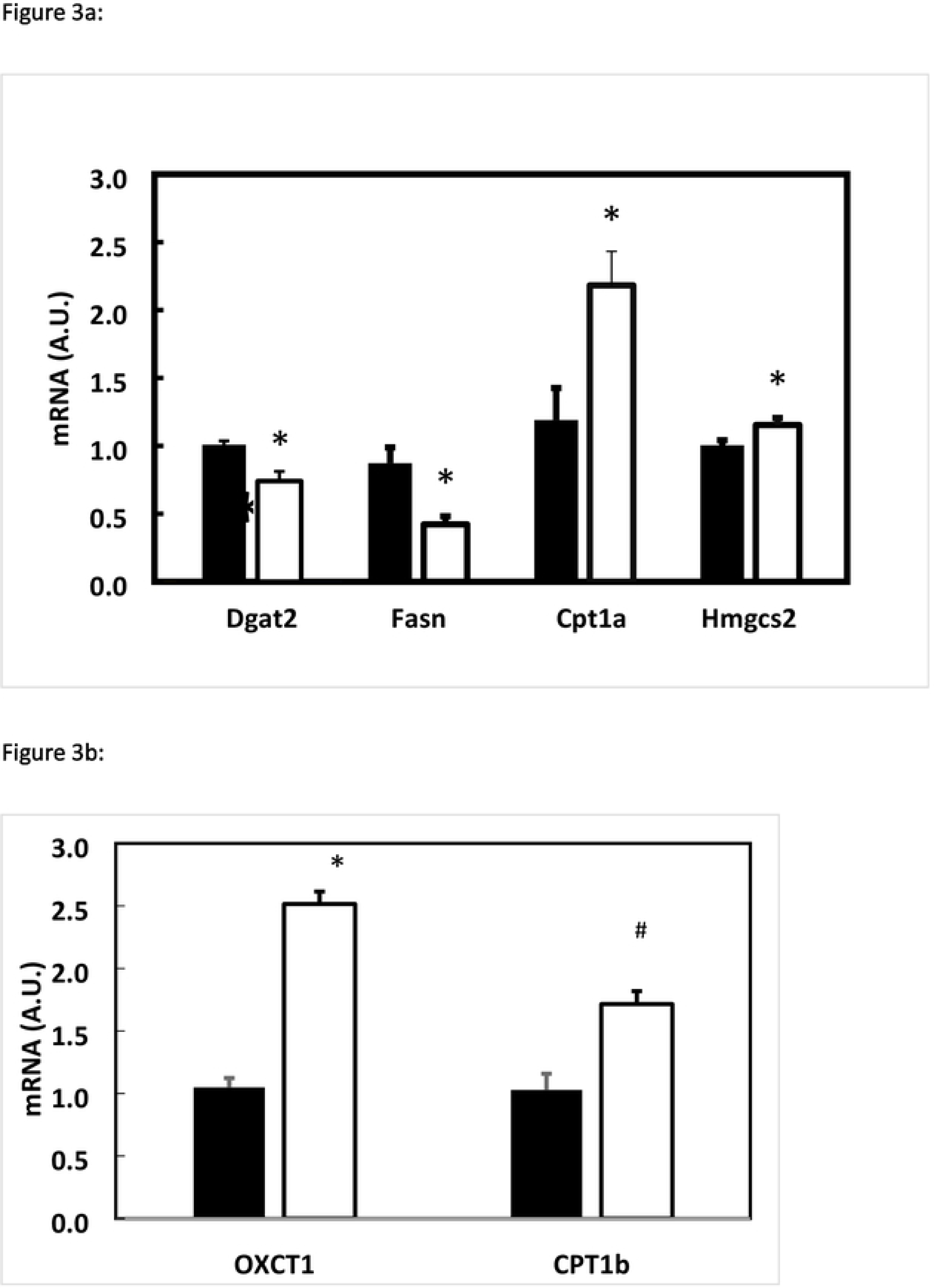
Effect of GSPE pre-treatment on gene expression in the short-challenge group at the end of the study. Rats were treated with 0.5 g/Kg BW for the first 10 days, and then they were put on a chow diet for 18 days and a cafeteria diet for 35 days. The black column indicates animals not treated with GSPE. The white column indicates animals treated with 0.5 g/Kg BW of GSPE for the first 10 days of treatment. The mRNA extracted from liver was quantified by RT-PCR and the relative gene expression detailed gens was obtained by the DDCt method in each gene. Figure 3a resumes liver results. Figure 3b summarizes muscle gene expression. The data are the mean ± SEM (n=7). Statistical differences identified by T-test are defined by * when p<0.05 between treatments.

Plasma ketone bodies were not significantly modified (control: **4.14** ± 0.35; GSPE: **3.89** ± 0.56; mM), so we measured the extent to which they could be oxidized by muscle. **Figure 3b** shows a strong significant increase in *Oxct1*(3-oxoacid CoA transferase 1) due to GSPE treatment, concomitantly with a tendency to increased C*pt1b*, which suggests that ketone bodies and fatty acids are the energy source in the muscle.

### Hormonal status of GSPE-treated rats after the short-cafeteria study

We next analysed the effects of GSPE on key hormones for the regulation of energetic homeostasis 7 weeks after finishing the GSPE treatment.

In pancreas, GSPE pre-treatment did not change insulin mRNA levels (**1.16** ± 0.25 in controls; **0.66** ± 0.11 in the GSPE pre-treated group; A.U.) but showed a tendency towards a lower glucagon mRNA (**1.16** ± 0.23; **0.60** ± 0.14; A. U.; p= 0.06). The amount of triglycerides(TG) in pancreas was not modified in GSPE pre-treated rats (**29.66** ± 2.21 in the control group and **27.10** ± 4.21 in the GSPE pre-treated group-μ g TG/ mg tissue).

GSPE pre-treatment statistically increased the plasma levels of total ghrelin (ng/mL; control: **4.09** ± 0.15; GSPE: **6.10** ± 0.42; p<0.05), although stomach mRNA levels of this hormone were not modified by GSPE pre-treatment (control: **1.04** ± 0.16; GSPE: **1.05** ± 0.12). In addition, GSPE pre-treatment led to a trend towards lower leptinemia (ng/mL; control: **28.0** ± 4.04; GSPE: **18.02** ± 2.15; p=0.07).

### Duration of GSPE effects

Finally, to estimate the duration of some of the effects described above, we analysed several parameters after a longer (17 weeks) cafeteria challenge.

The gene expression profile in liver differed considerably from that found in the short-challenge study (**table 3**). GSPE pre-treatment led to an increase in *Fasn* and *Dgat2* and a decrease in *Cpt1a* compared to the cafeteria treatment. This profile resembled that of the standard group of animals.

**Table 3.** Effects of GSPE on liver seventeen weeks after treatment (long cafeteria study) Liver samples were obtained at the end of the treatment. RT-PCR was used for each gene. The T test was used to determine statistical differences highlighted as *p<0.05 vs Cafeteria treatment; #: p<0.1 vs cafeteria group.

| Mesenteric WAT |  |  |
| --- | --- | --- |
| *Size of depot (g)* | $9.28 \pm 0.98$ | $5.96 \pm 0.62^*$ |
| *Cpt1b* | $1.05 \pm 0.08$ | $0.97 \pm 0.1$ |
| *Fasn* | $0.95 \pm 0.13$ | $0.62 \pm 0.1^{\#}$ |
| *Dgat2* | $1.01 \pm 0.09$ | $0.78 \pm 0.16$ |
| *Cpt1b/Fasn* | $1.22 \pm 0.27$ | $1.50 \pm 0.18$ |

The insulin/glucagon ratio of plasma levels in the GSPE pre-treated animals differed significantly from that of the cafeteria-fed animals, and produced a relationship closer to that of the standard-fed group (Chow: 0.86 ± 0.14*; cafeteria: 0.39 ± 0.10; GSPE: 0.95 ± 0.19*; *: p<0.05 vs cafeteria group).

## Discussion

Grape-seed derived proanthocyanidins have been extensively studied, but few studies use doses that are higher than can be provided by standard food ingestion but which may be interesting for a potential functional food. We showed that a dose of 0.5 g GSPE/kg bw has satiating properties in healthy rats^7^ and limits adipose accumulation induced by a cafeteria diet.^17^ We have also shown that GSPE maintains some of its antiobesogenic effects for a long period after GSPE administration finishes. In the present study, we analysed the mechanisms that explain this. We show that GSPE limits adipose fat pad accumulation until seven weeks after the last GSPE administration due to an inhibition in the adipose tissue *Lpl*. An increase in fatty acid oxidation in liver and muscle compensates for the inability of fatty acids to accumulate in WAT.

First, we show that GSPE also inhibits food intake if the diet is a tasty one (energy dense). During the ten-day GSPE treatment, animals reduced energy intake by 10% in comparison to the control group. Furthermore, these rats gained 30% less weight than the control (cafeteria) group.^17^ These results confirm that GSPE effects on food intake are additive to slimming effects because the lipid oxidation of GSPE is activated.^7^ In our short-term experiment, after GSPE intake, animals changed to a chow diet, and showed no changes in body weight accrual. Unexpectedly, when the rats were again fed a cafeteria diet (still without GSPE), the differences in body weight reappeared. These differences were around 40% between GSPE pre-treated animals and the control group, which correlate with the lower adiposity (79%),^17^ and there were no differences in either the quantity or quality of food intake. Our results, therefore, show that GSPE only has long-lasting anti-obesity effects under exposure to a high fat and/or a high sucrose diet.

A key element in triglyceride food-derived storage is the adipose lipoprotein lipase (*Lpl*). GSPE pre-treatment limited the amount of *Lpl* mRNA in mesenteric adipose tissue, which suggested an impairment of triglyceride storage in this tissue. *Lpl* has been shown to be a target for GSPE by Del Bas and col.^24^ They showed that five hours after an acute dose of 250 mg GSPE/kg bw there was a reduction in adipose *Lpl* mRNA. The results of the study by Yoon and col are more similar to ours.^25^ They showed that *Allomyrina dichotoma* larvae treatment had a considerable effect on *Lpl* mRNA because it limits adipose tissue growth in mice fed a high-fat diet. However, adipose *Lpl* limitation by itself is not sufficient to explain all GSPE effects. Weinstock and col, working with *Lpl* knockout mice that maintain *Lpl* expression only in muscle, showed no changes in the various adipose depots or in total lipid content.^26^ This suggests that in our study, GSPE might also act on other targets in the body.

If triglycerides cannot be stored in WAT after GSPE treatment, they might be derived to other organs. One of these organs is the liver, where GSPE pre-treatment directed fatty acids towards β-oxidation, as we found increased expression of CPT1a concomitantly with lower esterification (*Dgat2*). *Cpt1a* was also found to be up-regulated after two acute doses of 250 mg GSPE/kg bw^27^ in chow-fed rats but not after a subchronic treatment for 10 days with 25 mg/kg bw^5^ in 13-week cafeteria-fed rats. Baselga and col^28^ found a similar trend to ours in *Fasn* and *Cpt1a* mRNA levels after three weeks with a dose of 25 mg/kg bw in rats that had previously been on a cafeteria diet for 10 weeks. The main difference between our results and Baselga’s is that that they found that GSPE decreased liver triglycerides but we did not. On the contrary, our results show that triglycerides increased in the liver of the GSPE pre-treated rats.^29^ However, when we analyse the long cafeteria challenge (17 weeks) the triglyceride content in the GSPE pre-treated group did not differ from content in the vehicle-treated group, which suggests a limited trend toward their accumulation in liver. Yang-Xue and col,^30^ working with partially KO mice for *Lpl*, showed a strong *Lpl* mRNA inhibition in the youngest animals that was partially reduced with aging. These animals also showed an increased deposition of triglycerides in the liver in adulthood due to the KO gene that reverted the aging period. We worked with different treatments, different species and different durations, but we noticed a changing relationship between GSPE pre-treated rats and the amount of liver triglycerides, which suggests that time affects the accrual of triglycerides in the liver.

In the short-cafeteria study, GSPE pre-treated rats sent more triglycerides to the liver, oxidized fatty acids and produced ketone bodies, which were then removed by muscle. In fact, del Bas and col also showed an increase in the mRNA of muscle LPL,^24^ which suggests a derivation of fatty acids from adipose tissue to muscle that we cannot ignore. Similarly, fatty-acid uptake and oxidation were also found to be activated in muscle (higher mRNA *Cpt1b, Lpl* and *Cd36*) in males on a 10-week cafeteria diet that subsequently received 21 days of 25 mg GSPE/kg BW.^6^ Besides, the dose of GSPE does not seem to have a critical effect on muscle, as Crescenti and col found an overexpression of genes related to fatty acid uptake (Fatp1 and CD36) and b-oxidation in the skeletal muscle of STD-GSPE offsprings.^16^ So the higher oxidation of fatty acids in liver and muscle explain the lower RQ found in the GSPE pre-treated rats, a common trait of several GSPE treatments.^3^

It is important to point out that the metabolic changes remain for some considerable time after GSPE treatment. There may be several explanations for this. On the one hand, GSPE is an extract composed of absorbable compounds and non-absorbable structures. Absorbable compounds can reach the various tissues assayed ^31,32^ and non-absorbable structures can interact with intestinal sensors^33^. Through their interaction with microbiota, they can produce new compounds that can reach different targets in the body^34,35^. Thus, we cannot discount that there might be flavonoids remaining in the tissues. However, previous studies with a lower dose (100 mg GSPE/kg bw) administered for a longer time (12 weeks) suggested that flavonoids do not accumulate in liver or mesenteric adipose tissue.^36^ Another explanation might be that some epigenetic activity is taking place in the target tissues. GSPE modified liver miR-33a and miR-122^37^ at doses as low as 5 mg/kg BW for 3 weeks after a 15-week cafeteria diet. Similarly, Milenkovinc and col^38^ found changes in hepatic miRNA working with doses of proanthocyanidins closer to 5 mg/kg BW for two weeks. GSPE activity on histone deacetylases was proved by Downing and col^39^, who showed that GSPE regulates liver HDAC and Pparα activities to modulate lipid catabolism and reduce serum triglycerides *in vivo*. Similarly, Bladé and col proved that PACs modulate hepatic class III HDACs, which are often called sirtuins (SIRT1-7), in a dose-dependent manner. This was associated with significant protection against hepatic triglyceride and cholesterol accumulation in healthy rats.^10^ All this evidence, in conjunction with our findings on the regulation of gene expression in liver 7 weeks after GSPE treatment, suggests an epigenetic modelling of hepatic functioning by GSPE. The duration of these effects after GSPE administration is not clear. Crescenti and col showed effects in offspring 24 weeks after GSPE had been administered to their mothers during pregnancy.^40^ We observe that seventeen weeks after GSPE treatment *Ctp1a, Fasn* and *Dgat2* expression in the liver changed profile compared to 7 weeks after GSPE, which suggests that hepatic epigenetic regulation had come to an end. Instead, at this time point, the changes in liver clearly agree with the insulin/glucagon ratio modulation by GSPE. In fact, we showed that GSPE (45 days with 25 mg GSPE per kg of body weight) modulates pancreatic miRNAs.^41^ miR-483, which we showed is down-regulated by GSPE treatment in rat pancreatic islets, has been related to the optimum equilibrium between β-cells and α-cells (that is, its upregulation leads to increased insulin production and decreased glucagon synthesis).^42^ Therefore, our results also point to epigenetic changes in pancreas, which will need to be addressed in future work.

In conclusion, a short-term pre-treatment with GSPE repressed adipose *Lpl* and activated fatty oxidation in the liver. In conjunction with a greater utilization of ketone bodies in muscle, this would help to prevent an increase in body weight caused by a long-term high-fat diet after the end of treatment.

## Acknowledgements

We would like to thank Niurka Llopiz and the master student Marieke Schorstein for their technical support.

---

**Supplementary Figure 1. Schematic diagram of the experimental design**.

(1) CAF-LC: rats receiving a GSPE preventive treatment 10 days before the 17-week cafeteria intervention; (2) CAF-SC: rats receiving a GSPE preventive treatment for 10 days together with a high fat/high sucrose diet followed by an 18-day chow diet (standard diet) and then the 35-day cafeteria diet; GSPE: grape seed proanthocyanidin extract

## References

1. World Health Organization, World Heal. Organ. Media Cent. Fact Sheet No. 311, 2015.

2. E. P. Williams, M. Mesidor, K. Winters, P. M. Dubbert and S. B. Wyatt, Curr. Obes. Rep., 2015, 4, 363–370.

3. M. J. Salvadó, E. Casanova, A. Fernández-Iglesias, L. Arola and C. Bladé, Food Funct., 2015, 6, 1053–71.

4. S. Lamothe, N. Azimy, L. Bazinet, C. Couillard and M. Britten, Food Funct., 2014, 5, 2621–31.

5. H. Quesada, J. M. del Bas, D. Pajuelo, S. Díaz, J. Fernandez-Larrea, M. Pinent, L. Arola, M. J. Salvadó and C. Bladé, Int. J. Obes. (Lond)., 2009, 33, 1007–12.

6. E. Casanova, L. Baselga-Escudero, A. Ribas-Latre, L. Cedó, A. Arola-Arnal, M. Pinent, C. Bladé, L. Arola and M. J. Salvadó, J. Nutr. Biochem., 2014, 25, 1003–10.

7. J. Serrano, À. Casanova-Martí, A. Gual, A. M. A. M. Pérez-Vendrell, M. T. T. Blay, X. Terra, A. Ard?vol, M. Pinent, À. Casanova-Martí, A. Gual, A. M. A. M. Pérez-Vendrell, M. T. T. Blay, X. Terra, A. Ardévol and M. Pinent, Eur. J. Nutr., 2016, 56, 1629–1636.

8. J. Serrano, À. Casanova-Martí, I. Depoortere, M. T. M. T. Blay, X. Terra, M. Pinent and A. Ardévol, Mol. Nutr. Food Res., 2016, 60, 2554–2564.

9. K. Gil-Cardoso, I. Ginés, M. Pinent, A. Ardévol, L. Arola, M. Blay and X. Terra, Mol. Nutr. Food Res., 2017, 61, 1–12.

10. C. Bladé, G. Aragonès, A. Arola-Arnal, B. B. Muguerza, F. I. Bravo, M. J. Salvadó, L. Arola, M. Suárez, C. Blade, G. Aragones, A. Arola-Arnal, B. B. Muguerza, F. I. Bravo, M. J. Salvado, L. Arola and M. Suarez, Biofactors, 2016, 42, 5–12.

11. F. Puiggròs, N. Llópiz, A. Ardévol, C. Bladé, L. Arola, M. J. J. Salvadó, F. Puiggros, N. Llopiz, A. Ardevol, C. Blade, L. Arola, M. J. Salvado, N. Llópiz, A. Ardévol, C. Bladé, M. J. J. Salvadó, Ä. Piz, A. N. N. A. A. Rde, Ä. Vol and C. I. B. Lade, J. Agric. Food Chem., 2005, 53, 6080–6086.

12. N. Gonzalez-Abuin, M. Pinent, A. Casanova-Marti, L. Arola, M. Blay and A. Ardevol, Curr. Med. Chem., 2015, 22, 39–50.

13. X. Terra, G. Montagut, M. Bustos, N. Llopiz, A. Ardèvol, C. Bladé, J. Fernández-Larrea, G. Pujadas, J. Salvadó, L. Arola and M. Blay, J. Nutr. Biochem., doi:10.1016/j.jnutbio.2008.02.005.

14. A. Castell-Auvi, L. Cedo, V. Pallares, M. Blay, M. Pinent, A. Ardevol, A. Castell-Auví, L. Cedó, V. Pallarès, M. Blay, M. Pinent and A. Ardévol, J. Nutr. Biochem., 2013, 24, 948–953.

15. N. Gonzalez-Abuin, N. Martinez-Micaelo, M. Blay, A. Ardevol and M. Pinent, J. Agric. Food Chem., doi:10.1021/jf405239p.

16. A. Crescenti, J. M. del Bas, A. Arola-Arnal, G. Oms-Oliu, L. L. Arola and A. Caimari, J. Nutr. Biochem., 2015, 26, 912–920.

17. I. Ginés, K. Gil-Cardoso, J. Serrano, À. Casanova-Martí, Mt. Blay, I. Gin, K. Gil-Cardoso, J. Serrano, À. Casanova-mart and Mt. Blay, Nutrients, 2018, 10, 15.

18. M. Margalef, Z. Pons, L. Iglesias-Carres, L. Arola, B. Muguerza and A. Arola-Arnal, Mol. Nutr. Food Res., 2016, 60, 760–772.

19. J. Sánchez, T. Priego, M. Palou, A. Tobaruela, A. Palou, C. Picó, J. Sa, T. Priego, M. Palou, A. Tobaruela, A. Palou and C. Pico, Endocrinology, 2008, 149, 733–740.

20. B. P. Sampey, A. M. Vanhoose, H. M. Winfield, A. J. Freemerman, M. J. Muehlbauer, P. T. Fueger, C. B. Newgard and L. Makowski, Obesity, 2011, 19, 1109–1117.

21. J. Serrano, À. Casanova-Martí, K. Gil-Cardoso, M. T. Blay, X. Terra, M. Pinent and A. Ardévol, Food Funct., 2016, 7, 483–90.

22. A. Castell-Auvi, L. Cedo, V. Pallares, M. Blay, A. Ardevol and M. Pinent, Br. J. Nutr., 2012, 108, 1155–1162.

23. À. Casanova-martí, J. Serrano, K. J. Portune and Y. Sanz, Food Funct., doi:10.1039/c7fo02028g.

24. J. M. M. Del Bas, J. Fernández-larrea, M. Blay, A. Ardèvol, M. J. M. J. Salvadó, L. Arola, C. Bladé, J. Fernandez-Larrea, M. Blay, A. Ardevol, M. J. Salvado, L. Arola, C. Blade, J. Maria, D. Bas, J. Fernández-larrea, M. Blay, A. Ardèvol, M. J. M. J. Salvadó, J. M. M. Del Bas, J. Fernández-larrea, M. Blay, A. Ardèvol, M. J. M. J. Salvadó, L. Arola, C. Bladé, J. Fernandez-Larrea, M. Blay, A. Ardevol, M. J. Salvado, L. Arola and C. Blade, FASEB J., 2005, 19, 479–81.

25. S. K. Fried, C. D. Russell, N. L. Grauso and R. E. Brolin, Insulin, 1993, 92, 2191–2198.

26. P. H. Weinstock, C. L. Bisgaier, K. Aalto-Set„l„, H. Radner, R. Ramakrishnan, S. Levak-Frank, A. D. Essenburg, R. Zechner and J. L. Breslow, J. Clin. Invest., 1995, 96, 2555–2568.

27. J. M. J. M. Del Bas, M. L. M. L. Ricketts, I. Baiges, H. Quesada, A. Ardevol, M. J. Salvado, G. Pujadas, M. Blay, L. Arola, C. Blade, D. D. D. Moore, J. Fernandez-Larrea, J. Maria, D. Bas, M. L. M. L. Ricketts, I. Baiges, H. Quesada, A. Ardevol, M. J. M. J. Salvadó, G. Pujadas, M. Blay, L. Arola, C. Bladø, D. D. D. Moore, J. Fernandez-Larrea, C. Bladé, D. D. D. Moore and J. Fernandez-Larrea, Mol. Nutr. Food Res., 2008, 52, 1172–1181.

28. L. Baselga-Escudero, A. Arola-Arnal, A. Pascual-Serrano, A. Ribas-Latre, E. Casanova, M.-J. J. Salvadó, L. Arola and C. Blade, PLoS One, 2013, 8, 1–8.

29. Y.-I. Yoon, M. Chung, J.-S. Hwang, M. Han, T.-W. Goo and E.-Y. Yun, Nutrients, 2015, 7, 1978–1991.

30. Y. X. Li, T. T. Han, Y. Liu, S. Zheng, Y. Zhang, W. Liu and Y. M. Hu, Atherosclerosis, 2015, 239, 276–282.

31. A. Ardévol, M. J. J. Motilva, A. Serra, M. Blay, M. Pinent, A. Ardevol, M. J. J. Motilva, A. Serra, M. Blay, M. Pinent, A. Ardévol, M. J. J. Motilva, A. Serra, M. Blay and M. Pinent, Food Chem., 2013, 141, 160–6.

32. M. Margalef, Z. Pons, F. I. Bravo, B. Muguerza and A. Arola-Arnal, J. Nutr. Biochem., 2015, 26, 987–995.

33. W. S. U. Roland, R. J. Gouka, H. Gruppen, M. Driesse, L. van Buren, G. Smit and J.-P. Vincken, PLoS One, 2014, 9, 1–10.

34. A. Casanova-Marti, Food Funct., 2018, 9, 1672–1682.

35. M. Margalef, Z. Pons, F. I. Bravo, B. Muguerza and A. Arola-Arnal, J. Funct. Foods, 2015, 12, 478–488.

36. M. Margalef, Z. Pons, L. Iglesias-Carres, F. I. Bravo, B. Muguerza and A. Arola-Arnal, J. Agric. Food Chem., 2015, 63, 9996–10003.

37. L. Baselga-Escudero, A. Pascual-Serrano, A. Ribas-Latre, E. Casanova, M. J. Salvadó, L. Arola, A. Arola-Arnal and C. Bladé, Nutr. Res., 2015, 35, 337–345.

38. D. Milenkovic, C. Deval, E. Gouranton, J. F. Landrier, A. Scalbert, C. Morand and A. Mazur, PLoS One, doi:10.1371/journal.pone.0029837.

39. L. E. Downing, B. S. Ferguson, K. Rodriguez and M.-L. Ricketts, Mol. Nutr. Food Res., 2017, 61, 1600347.

40. J. M. del Bas, A. Crescenti, A. Arola-Arnal, G. Oms-Oliu, L. Arola and A. Caimari, Int. J. Obes. (Lond)., 2015, 39, 7–15.

41. A. Castell-Auvi, L. Cedo, J. Movassat, B. Portha, F. Sanchez-Cabo, V. Pallares, M. Blay, M. Pinent, A. Ardevol, A. Castell-Auví, L. Cedó, J. Movassat, B. Portha, F. Sánchez-Cabo, V. Pallarès, M. Blay, M. Pinent, A. Ardévol and A. Arde, J. Agric. Food Chem., 2013, 61, 355–363.

42. R. Mohan, Y. Mao, S. Zhang, Y. W. Zhang, C. R. Xu, G. Gradwohl and X. Tang, J. Biol. Chem., 2015, 290, 19955–19966.

